# A nonlinear updating algorithm captures suboptimal inference in the presence of signal-dependent noise

**DOI:** 10.1101/258434

**Authors:** Seth W. Egger, Mehrdad Jazayeri

## Abstract

Bayesian models of behavior have advanced the idea that humans combine prior beliefs and sensory observations to minimize uncertainty. How the brain implements Bayes-optimal inference, however, remains poorly understood. Simple behavioral tasks suggest that the brain can flexibly represent and manipulate probability distributions. An alternative view is that brain relies on simple algorithms that can implement Bayes-optimal behavior only when the computational demands are low. To distinguish between these alternatives, we devised a task in which Bayes-optimal performance could not be matched by simple algorithms. We asked subjects to estimate and reproduce a time interval by combining prior information with one or two sequential measurements. In the domain of time, measurement noise increases with duration. This property makes the integration of multiple measurements beyond the reach of simple algorithms. We found that subjects were able to update their estimates using the second measurement but their performance was suboptimal, suggesting that they were unable to update full probability distributions. Instead, subjects’ behavior was consistent with an algorithm that predicts upcoming sensory signals, and applies a nonlinear function to errors in prediction to update estimates. These results indicate that inference strategies humans deploy may deviate from Bayes-optimal integration when the computational demands are high.

## Introduction

Sensorimotor control depends on accurate estimation of internal state variables [1–5]. Numerous experiments have used Bayesian estimation theory to demonstrate that humans estimate internal states by integrating multiple sources of information including prior beliefs and sensory cues from various modalities [6–16]. Bayesian estimation is typically formulated in terms of three components: prior distributions representing *a priori* beliefs about state variables, likelihood functions derived from noisy sensory measurements, and cost functions that characterize reward contingencies [17]. In this formulation, the likelihood function and prior distribution are combined to compute a posterior distribution and the cost function is used to extract an estimate that maximizes expected reward. This formulation is the basis of most psychophysical studies of Bayesian integration [9–15, 18–20].

Implicit in this formulation is the assumption that the brain has access to priors, likelihoods, and cost functions. Access to these quantities is appealing as it could support rapid and optimal state estimation without the need to learn new policies for novel behavioral contexts [21, 22]. However, in most experiments, Bayes-optimal behavior can also be achieved by simpler algorithms that do not depend on direct access to likelihoods, priors and cost functions [21–23]. For example, optimal cue combination in the presence of Gaussian noise may be implemented by a weighted sum of measurements [6]. Similarly, integration of noisy evidence with prior beliefs may be implemented by a suitable functional mapping between measurements and estimates [24, 25]. Finally, online estimation of a variable from sequential measurements that are subject to Gaussian noise can be achieved by a *Kalman filter* that only keeps track of the mean and variance [26] without representing and updating the full posterior distribution.

In contrast to simple laboratory tasks, optimal inference in natural settings is often intractable and involves approximations that may deviate from optimality [27]. Therefore, it is critical to go beyond statements of optimality and suboptimality, and assess the inference algorithms humans use during sensorimotor and cognitive tasks [28, 29]. Further, as articulated by Marr [30], characterization of the underlying algorithms could establish a bridge between behaviorally relevant computations and neurobiological mechanisms.

We devised an experiment in which the computational demands for optimal Bayesian estimation were incompatible with simple algorithmic solutions. Subjects had to reproduce an interval by integrating their prior belief with one or two measurements of the interval. In the domain of time, the noise is signal-dependent, which causes behavioral variability to scale with duration [31]. Theoretical considerations suggest that an important consequence of scalar variability is that simple algorithms that only update certain parameters of the posterior (e.g., mean and/or variance) cannot emulate Bayes-optimal behavior. Therefore, optimal behavior in this paradigm would provide strong evidence that the underlying inference algorithm involves updating probability distributions. Conversely, suboptimal behavior would suggest that subjects rely on a simpler algorithm. We found that when subjects made two measurements their performance was suboptimal. Furthermore, comparison of behavior with various models indicated that subjects relied on an inference algorithm that used measurements to update point estimates using point nonlinearities.

## Results

### Subjects integrate interval measurements with prior knowledge

Subjects performed an interval reproduction task consisting of two randomly interleaved trial types (Fig 1A,B). In “1-2-Go” trials, two flashes (S1 followed by S2) demarcated a sample interval (*t_s_*). Subjects had to reproduce *t_s_* immediately after S2. The time between the onset of S2 and when the keyboard was pressed was designated as the production interval (*t_p_*). In “1-2-3-Go” trials, *t_s_* was presented twice, demarcated once by S1 and S2 and once by S2 and S3, providing the opportunity to make two measurements (Fig 1B). Similar to 1-2-Go, subjects had to match *t_p_* (the interval between S3 and keyboard press) to *t_s_*. Across trials, *t_s_* was drawn from a discrete uniform distribution ranging between 600 and 1000 ms (Fig 1C). Subjects received two forms of trial-by-trial feedback based on the magnitude and sign of the error. First, a feedback stimulus was presented whose location relative to the warning stimulus reflected the magnitude and sign of the error (Fig 1A,B; see Materials and methods). Second, if the error exceeded a threshold window (Fig 1D), stimuli remained white and a tone denoting incorrect response was presented. Otherwise, the stimuli turned green and a tone denoting correct was presented. The threshold window for correct performance was proportionally larger for the longer *t_s_* to accommodate the scalar variability of timing due to signal-dependent noise [40–45]. The threshold was adjusted adaptively and on a trial-by-trial basis according to performance (see Materials and methods).

**Fig 1.**
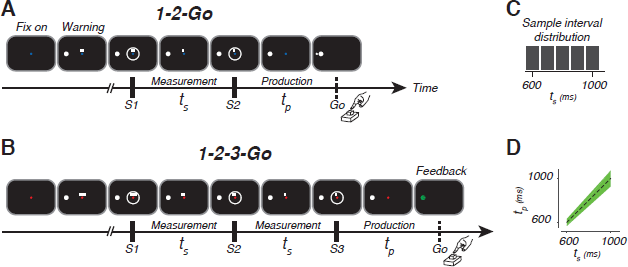
The 1-2-Go and 1-2-3-Go interval reproduction task. (A,B) Task design. Each trial began with the appearance of a fixation spot (Fix on). The color of the fixation spot informed the subject of the trial type: blue for 1-2-Go, and red for 1-2-3-Go. After a random delay, a warning stimulus (large white circle) appeared. Additionally, two or three small white rectangles were presented above the fixation spot. The number of rectangles was associated with the number of upcoming flashes. After another random delay, two (S1 and S2 for 1-2-Go) or three (S1, S2 and S3 for 1-2-3-Go) white annulae were flashed for 100 ms in a sequence around the fixation spot. Consecutive flashes were separated by the duration of the sample interval (*t_s_*). With the disappearance of each flash, one of the small rectangles also disappeared (rightmost first and leftmost last). The white rectangles were provided to help subjects keep track of events during the trial. Subjects had press a button after the last flash to produce an interval (*t_p_*) that matched *t_s_*. Immediately after button press, subjects received analog feedback. The analog feedback was a small circle that was presented to the left or right of the warning stimulus depending on whether *t_p_* was larger or smaller than *t_s_*, respectively. The distance of the feedback circle to the center of the warning stimulus was proportional to the magnitude of the error (*t_p_ − t_s_*). (C) Experimental distribution of sample intervals. (D) Feedback. Subjects were given positive feedback if their production times fell within the green region. The width of the positive feedback window was scaled with *t_s_*.

Subjects’ timing behavior exhibited three characteristic features (Fig 2). First, *t_p_* increased monotonically with *t_s_*. Second, *t_p_* was systematically biased toward the mean of the prior, as evident from the tendency of responses to deviate from *t_s_* (diagonal) and gravitate toward the mean *t_s_*. As proposed previously [24, 34–36], this so-called regression to the mean indicated that subjects relied on their knowledge of the prior distribution of *t_s_*. Third, performance was better in 1-2-3-Go condition in which subjects made two measurements, as evidenced by a lower root-mean-square error (RMSE) in 1-2-3-Go compared to 1-2-Go condition (Fig 2C; permutation test; p-value *<* 0.01 for all subjects). This observation indicates that subjects combined the two measurements to improve their estimates, corroborating reports from other behavioral paradigms [46–54]. Combined with the systematic bias toward the mean of *t_s_*, these results indicated that subjects integrated prior information with one or two measurements to improve their performance.

### A Bayesian model of behavior

Building Fig 2. Performance in the interval on previous work [11, 18, 22, 55], we asked whether subjects’ behavior could be accounted for by a Bayesian observer model based on the Bayes-Least Squares (BLS) estimator. For the 1-2-Go trials, the observer model (1) makes a noisy measurement of *t_s_*, which we denote by *t_m_*_1_, (2) combines the likelihood function associated with *t_m_*_1_, *p*(*t_m_*_1_*|t_s_*), with the prior distribution of *t_s_*, *p*(*t_s_*), to compute the posterior, *p*(*t_s_|t_m_*_1_), and (3) uses the mean of the posterior as the optimal estimate, *t_e_*_1_. We modeled *p*(*t_m_*_1_*|t_s_*) as a Gaussian distribution centered at *t_s_* with standard deviation, *σ_m_*, proportional to *t_s_* with constant of proportionality, *w_m_*; i.e., *σ_m_* = *w_m_t_s_* (Fig 3A, left box). We assumed that the production process was also perturbed by noise and modeled *t_p_* as a sample from a Gaussian distribution centered at *t_e_*_1_ with standard deviation, *σ_p_*, proportional to *t_e_*_1_ with constant of proportionality, *w_p_*; i.e., *σ_p_* = *w_p_t_e_*_1_ (Fig 3A, right box). Note that the entire operation of the BLS estimator can be described in terms of a deterministic mapping of *t_m_*_1_ to *t_e_*_1_ using a nonlinear function, which we denote as *f_BLS_*_1_(*t_m_*_1_) (Fig 3B) [24].

**Fig 2.**
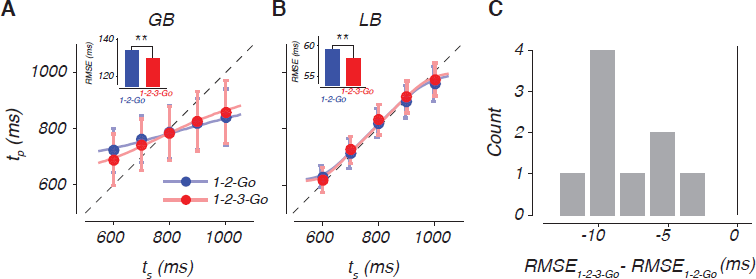
Performance in the interval reproduction task. (A) Production interval (*t_p_*) as a function of sample interval (*t_s_*) for a low sensitivity subject. Filled circles and error bars show the mean and standard deviation of *t_p_* for each *t_s_* in the 1-2-Go (blue) and 1-2-3-Go (red) conditions. The dotted unity line represents perfect performance and the colored lines show the expected *t_p_* from a Bayes Least-Squares (BLS) model fit to the data. Inset: root-mean-square error (RMSE) in the 1-2-Go (blue) and 1-2-3-Go (red) conditions differed significantly (asterisk, p-value *<* 0.01; permutation test). (B) Same as (A) for a high sensitivity subject (LB). (C) The histogram of changes in RMSE across conditions for all subjects.

For the 1-2-3-Go trials, the subject makes two measurements, *t_m_*_1_ and *t_m_*_2_, and uses the mean of the posterior based on both measurements, *p*(*t_s_|t_m_*_1_*, t_m_*_1_), to derive an optimal estimate, *t_e_*_2_ (Fig 3C; see Materials and methods). In these trials, the mapping from *t_m_*_1_ and *t_m_*_2_ to *t_e_*_2_ can be described in terms of a two-dimensional nonlinear function, denoted by *f_BLS_*_2_(*t_m_*_1_*, t_m_*_2_) (Fig. 3D). Note that the iso-estimate contours of *f_BLS_*_2_(*t_m_*_1_*, t_m_*_2_) are both nonlinear and convex (Fig. 3D, red). The nonlinearity indicates that the effect of *t_m_*_1_ and *t_m_*_2_ on *t_e_*_2_ is non-separable, and the convexity indicates that *t_e_*_2_ is more strongly influenced by the larger of the two measurements. These features are direct consequences of scalar noise and are not present when measurements are perturbed by Gaussian noise (see S1 Appendix).

We fit the model to each subject’s data assuming that responses in both 1-2-Go and 1-2-3-Go conditions were associated with the same *w_m_* and *w_p_* (Materials and methods). The model was augmented in two ways to ensure that estimates of *w_m_* and *w_p_* were accurate. First, we included an offset parameter to absorb interval-independent biases (e.g., consistently pressing the button too early or too late). Second, trials in which *t_p_* grossly deviated from *t_s_* were designated as “lapse” trials (see Materials and methods).

**Fig 3.**
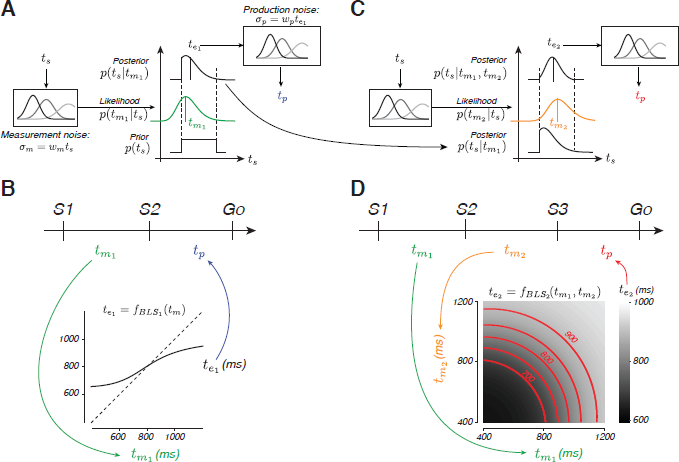
BLS model of interval integration. (A) BLS model for 1-2-Go trials. The left panel illustrates the measurement process. The measured interval, *t_m_*_1_, is perturbed by zero-mean Gaussian noise whose standard deviation is proportional to the sample interval, *t_s_*, with constant of proportionality *w_m_* (*σ_m_* = *w_m_t_s_*). The middle panel illustrates the estimation process. The model multiplies the likelihood function associated with *t_m_*_1_ (middle panel, green) with the prior (bottom), and uses the mean of the posterior (top) to derive an interval estimate (*t_e_*_1_, black vertical line on the posterior). The right panel illustrates the production process. The produced interval, *t_p_*, is perturbed by zero-mean Gaussian noise with standard deviation proportional to *t_e_*_1_, with constant of proportionality *w_p_* (*σ_p_* = *w_p_t_e_*_1_). (B) The effective mapping function (*f_BLS_*_1_, black curve) from the first measurement, *t_m_*_1_, to the optimal estimate, *t_e_*_1_. The dashed line indicates unity. (C) BLS model for 1-2-3-Go trials. The model uses the posterior after the first measurement, *p*(*t_s_|t_m_*_1_), as the prior and combines it with the likelihood of the second measurement (*t_m_*_2_, orange) to compute an updated posterior, *p*(*t_s_|t_m_*_1_*, t_m_*_2_). The mean of the updated posterior is taken as the interval estimate (*t_e_*_2_). (D) The effective mapping function (*f_BLS_*_2_, grayscale) from each combination of measurements, *t_m_*_1_ and *t_m_*_2_, to the optimal the estimate, *t_e_*_2_. Red lines indicate combinations of measurements that lead to identical estimates (shown for *t_e_*_2_ = 700, 750, 800, 850, and 900 ms).

Model fits captured subjects’ behavior for both conditions as shown by a few representative subjects (Fig 4A, 2A and 2B; see Supporting information for fits to all the subjects). Following previous work [24], we evaluated model fits using two statistics, an overall bias, BIAS, and an overall variability, 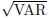 (see Materials and methods). As shown in Fig 4B, the model broadly captured the bias and variance for all subjects in both 1-2-Go and 1-2-3-Go conditions. However, the observed BIAS was significantly larger than predicted by the model fits in the 1-2-3-Go condition (Fig 4B, inset; two tailed t-test, t(8) = 4.6982, p-value = 0.0015), but not in the 1-2-Go condition (two tailed t-test, t(8) = −0.3236, p-value = 0.7546).

We quantified this observation across subjects by normalizing each subject’s RMSE in the 1-2-Go and 1-2-3-Go conditions to the RMSE expected from the BLS model in the 1-2-3-Go condition (Fig 4C). We found that the observed RMSE in the 1-2-3-Go condition was significantly larger than expected (two tailed t-test, t(8) = 3.5484, p-value = 0.007). Further, the drop in observed RMSE in the 1-2-3-Go was significantly less than expected by the BLS model (one tailed t-test, t(8) = −4.4600, p-value = 0.0011). These analyses indicate that subjects were able to integrate the two measurements but failed to optimally update the posterior by the likelihood information associated with the second measurement.

**Fig 4.**
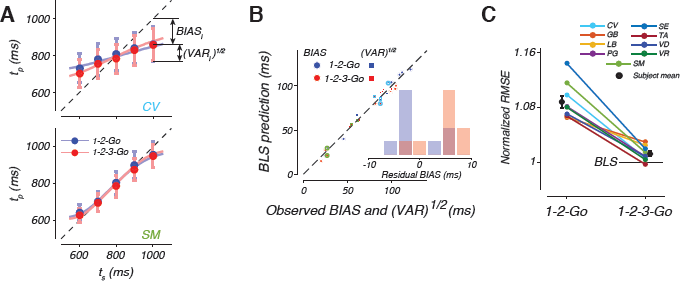
BLS model fits to data. (A) Behavior of two subjects and the corresponding BLS model fits with the same format as in Fig 2A and 2B. (B) BIAS (circles) and 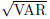 (squares) of each subject (abscissa) and the corresponding values computed from simulations of the fitted BLS model (ordinate). Red and blue points correspond to 1-2-Go and 1-2-3-Go, respectively. The dotted line plots unity. Data points corresponding to subjects SM and CV are marked by light green and light blue, respectively. Inset: difference between the BIAS observed from data and that expected by the BLS model fit for 1-2-Go (blue) and 1-2-3-Go (red) conditions. (C) Comparison of behavioral performance to model predictions. Each line connects the RMSE for 1-2-Go (left) and 1-2-3-Go (right) conditions for one subject. To facilitate comparison across subjects, RMSE values for each subject were normalized by the RMSE of the BLS model in the 1-2-3-Go condition. The black circles and error bars correspond to the mean and standard error of the normalized RMSE across subjects. See also Supporting information.

### An algorithmic view of Bayesian integration

The success of the BLS model in capturing behavior in the 1-2-Go condition [24, 34, 36] and its failure in the 1-2-3-Go condition suggests that subjects were unable to update the posterior by the second measurement. We examined a number of simple inference algorithms that could account for this limitation. One of the simplest algorithms proposed for integrating sequential measurements is the Kalman filter. The Kalman filter only updates the mean and variance of the posterior [26]. This strategy is optimal when measurement noise is Gaussian because a Gaussian distribution is fully determined by its mean and variance. More generally, when integrating the likelihood function leaves the parametric form of the posterior distribution unchanged, a simple inference algorithm that updates those parameters can implement optimal integration.

First, we asked whether there exists a similarly simple and optimal updating algorithm when the noise is signal-dependent (i.e., scalar noise). For the posterior to have the same parametric form after one and two measurements, it is necessary that the product of two likelihood functions have the same parametric form as a single likelihood function. We tested this property analytically and verified that the parametric form of the likelihood function associated with scalar noise was not invariant under multiplication (see S1 Appendix). Therefore, any inference algorithm that only updates certain statistics of the posterior (e.g., mean and variance) is expected to behave suboptimally when multiple time intervals have to be integrated. Therefore, we hypothesized that subjects might have used a simple updating algorithm analogous to the Kalman filter to integrate multiple measurements.

### A linear-nonlinear estimator (LNE) model for approximate Bayesian inference

The first algorithm we tested was one in which the observer combines the last estimate *t_n−_*_1_, with the current measurement, *t_m__n_*, using a linear updating strategy. If we denote the corresponding weights by 1 *− k_n_* and *k_n_* and set *k_n_* = 1*/n*, this algorithm tracks the running average of the measurements, *t_n_* (*k*_1_ and *k*_2_ are 1 and 0.5, respectively). However, such linear updating scheme would certainly fail to account for the observed nonlinearities in subjects’ behavior (S2 Fig). Therefore, we constructed a linear-nonlinear estimator (LNE) that augmented the linear updating by a point nonlinearity that could account for the observed prior dependent biases in *t_p_* (Fig 5A). The nonlinear function, *f_BLS_*_1_(*t_n_*), was chosen to match the BLS estimator for a single measurement (*n* = 1), which is determined by *w_m_*.

Simulation of LNE verified that it could indeed integrate multiple measurements and exhibit prior-dependent biases (not shown). However, the behavior of LNE was qualitatively different from BLS. The contrast between the two models was evident from a comparison of the relationship between measurements and estimates. Unlike BLS (Fig 3D), estimates derived from LNE are linear with respect to *t_m_*_1_ and *t_m_*_2_, a feature that can be visualized by the linear iso-estimate contours of the LNE model (Fig 5B).

We fitted LNE to each subject independently and asked how well it accounted for the observed statistics. The LNE model broadly captured the observed regression to the mean (Fig 5C,D; see S4 Fig for fits to all subjects), but had a qualitative failure: fits exhibited significantly more BIAS in 1-2-Go condition (Fig 5D, inset; two tailed t-test, t(8) = 4.9304, p-value = 0.001) and significantly less BIAS in 1-2-3-Go condition (Fig 5D, inset; two tailed t-test, t(8) = −2.3782, p-value = 0.045) than the biases present in the data. This failure can be readily explained in terms of how LNE functions. Since the static nonlinearity in LNE is the same for one and two measurements, the bias LNE generates is the same for the 1-2-Go and 1-2-3-Go conditions. Therefore, when we fitted LNE to data from both conditions, the model consistently overestimated BIAS for the 1-2-Go condition, and underestimated BIAS for the 1-2-3-Go condition (Fig 5C, red and blue lines nearly overlap).

We further evaluated LNE by asking how it accounted for the observed performance improvement in the 1-2-3-Go condition compared to the 1-2-Go condition. We normalized each subject’s RMSE from the 1-2-Go and 1-2-3-Go conditions to the RMSE expected from the behavior of the fitted LNE model in the 1-2-3-Go condition (Fig 5E). Most subjects surpassed the predictions of the LNE model (horizontal line) for the 1-2-3-Go condition, and average RMSE reached values that were significantly lower than expected (0.990; two tailed t-test, t(8) = −2.463, p-value = 0.039). Based on these results, we concluded that LNE fails to capture subjects’ behavior both qualitatively and quantitatively.

### An extended Kalman filter (EKF) model for approximate Bayesian inference

We considered a moderately more sophisticated algorithm inspired by the extended Kalman filter (EKF) [56]. This algorithm is shown in Fig 6A. Upon each new measurement, EKF uses the error between the previous estimate and the current measurement to generate a new estimate. The difference between EKF and the Kalman filter is that the error is subjected to a nonlinear function before being used to update the previous estimate. This nonlinearity is necessary for the algorithm to be able to account for the nonlinear prior-dependent biases observed in behavior.

**Fig 5.**
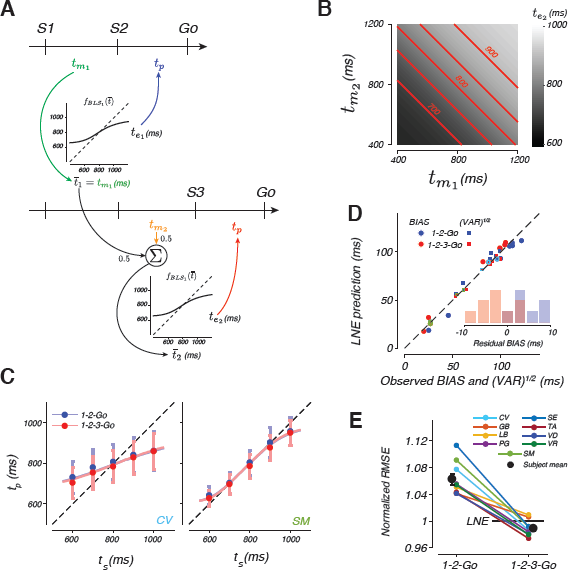
A linear-nonlinear estimator (LNE) model and its fits to the data. (A) LNE algorithm. LNE derives an estimate by applying a nonlinear function, *f_BLS_*_1_, to the average of the measurements. In the 1-2-Go trials (top), the average, ¯*t*_1_, is the same as the first measurement, *t_m_*_1_, and the estimate, *t_e_*_1_, is *f_BLS_*_1_(¯*t*_1_). In 1-2-3-Go trials (bottom), the average, ¯*t*_2_, is updated by the second measurement, *t_m_*_2_ (¯*t*_2_ = 0.5(¯*t*_1_ C *t_m_*_2_)), and the estimate, *t_e_*_2_, is *f_BLS_*_1_(¯*t*_2_). In both conditions, the produced interval, *t_p_*, is perturbed by zero-mean Gaussian noise with standard deviation proportional to the final estimate (*t_e_*_1_ for 1-2-Go and *t_e_*_2_ for 1-2-3-Go) with the constant of proportionality *w_p_*, as in the BLS model. (B) The mapping from measurements to estimates (grayscale) for the LNE estimator in the 1-2-3-Go trials. Red lines indicate combinations of measurements that lead to identical estimates (shown for *t_e_*_2_ = 700, 750, 800, 850, and 900 ms). (C) Mean and standard deviation of *t_p_* as a function of *t_s_* for two example subjects (circles and error bars) along with the corresponding fits of the LNE model (lines). (D) BIAS (circles) and 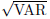 (squares) of each subject (abscissa) and the corresponding values computed from simulations of the fitted LNE model (ordinate). Conventions match Fig 4B. (E) The RMSE in the 1-2-Go and 1-2-3-Go conditions relative to the corresponding predictions from the LNE model (conventions as in Fig 4C). See also S4 Fig.

In our experiment, immediately after the first flash, the only information about the sample interval, *t_s_*, comes from the prior distribution. Accordingly, we set the initial estimate, *t_e_*_0_, to the mean of the prior distribution. After the first measurement, EKF computes an “innovation” term by applying a static nonlinearity, *f^*^*(*x*) to the error, *x*_1_ between *t_m_*_1_ and *t_e_*_0_. This innovation is multiplied by a gain, *k*_1_ and added to *t_e_*_0_ to compute the new estimate, *t_e_*_1_. In the 1-2-Go condition in which only one measurement is available, *t_e_*_1_ serves as the final estimate that the model aims to reproduce.

For the 1-2-3-Go condition, EKF repeats the updating procedure after the second measurement *t_m_*_2_. It computes the difference between *t_m_*_2_ and *t_e_*_1_ to derive a prediction error, *x*_2_, which is subjected to the same nonlinear function, *f^*^*(*x*), to yield a second innovation. This innovation is then scaled by an appropriate gain, *k*_2_, and added to *t_e_*_1_ to generate an updated estimate, *t_e_*_2_, which the model aims to reproduce.

The two important elements that determine the overall behavior of EKF are the nonlinear function *f^*^*(*x*) and the gain factor(s) applied to the innovation(s) (*k*_1_ and *k*_2_) to update the estimate(s). We set the form of the nonlinear function *f^*^*(*x*) such that biases in *t_e_*_1_ after one measurement are the same between EKF and BLS models. This ensures that EKF and BLS behave identically in the 1-2-Go condition. Note that our implementation of EKF assumes that the same nonlinear function is applied after every measurement. If one allows this nonlinear function to be optimized separately for each measurement, EKF would be able to replicate the behavior of BLS exactly (S3 Fig).

For the gain factors, we reasoned that the most rational choice is to set the weight of each innovation based on the expected reliability of the corresponding estimate, *t_e__n−_*_1_, relative to the new measurement, *t_m__n_*, as in the Kalman filter (see Materials and methods). This causes the gain factor to decrease with the number of measurements, and ensures that the influence of each new measurement is appropriately titrated. With these assumptions, EKF remains suboptimal for the 1-2-3-Go condition. However, it captures certain aspects of the nonlinearities associated with the optimal BLS estimator as shown by Fig 6B (compare to Fig 3D).

The algorithm implemented by EKF is appealing as it uses a simple updating strategy that can be straightforwardly extended to multiple sequential measurements. Furthermore, EKF captures important features of human behavior. First, integration of each new measurement causes a reduction in RMSE, as seen in 1-2-3-Go compared to 1-2-Go condition. Second, the nonlinear function applied to innovations allows EKF to incorporate prior information and capture prior-dependent biases. Third, since the nonlinearity is applied to each innovation (as opposed to the final estimate), EKF, unlike LNE, is able to capture the reduction in BIAS in 1-2-3-Go compared to 1-2-Go condition.

We fitted EKF to each subject independently and asked how well it accounted for the observed statistics. Similar to BLS and LNE, EKF broadly captured the observed regression to the mean in the 1-2-Go trials (Fig 6C,D, blue). This is not surprising since the EKF algorithm is identical to BLS when the prior is integrated with a single measurement. EKF was also able to capture the mean *t_p_* as a function of *t_s_* in the 1-2-3-Go trials (Fig 6C,D red). Importantly, unlike BLS, there was no significant difference between the BIAS observed in subject behavior and the EKF model (Fig 6D, inset; two tailed t-test, t(8) = 4.6055, p-value = 0.02639).

We also asked if EKF could account for the observed RMSEs. To do so, we performed the same analysis we used to evaluate the BLS and LNE models. We normalized each subject’s RMSE from the 1-2-Go and 1-2-3-Go conditions to the RMSE expected from the EKF model for 1-2-3-Go (Fig 6E). We found no significant difference between observed and predicted RMSEs for the 1-2-3-Go condition (two-tailed t-test, t(8) = 1.5506, p-value = 0.160), and no significant difference between the observed and predicted change in RMSE from the 1-2-Go to the 1-2-3-Go condition (one tailed t-test, t(8) = −1.1090, p-value = 0.150). These results indicate that subjects’ suboptimal behavior is consistent with the approximate Bayesian integration implemented by the EKF algorithm.

**Fig 6.**
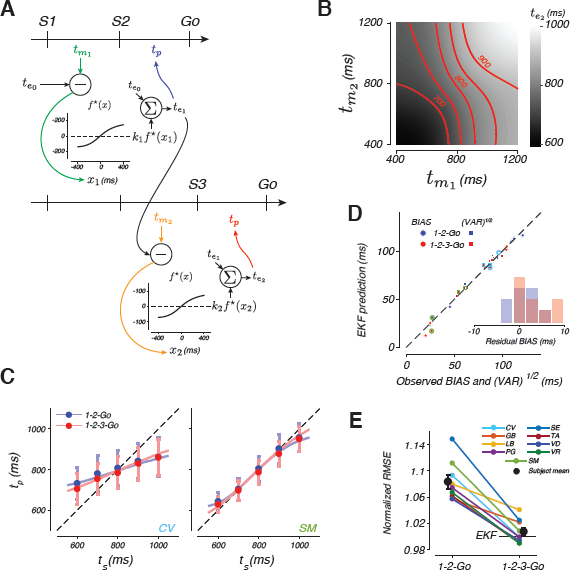
An extended Kalman filter (EKF) model and its fits to the data. (A) EKF algorithm. EKF is a real-time inference algorithm that uses each measurement to update the estimate. After the first flash, EKF uses the mean of the prior as its initial estimate, *t_e_*_0_. The second flash furnishes the first measurement, *t_m_*_1_. EKF computes a new estimate, *t_e_*_1_, using the following procedure: (1) it measures the difference between *t_m_*_1_ and *t_e_*_0_ to compute an error, *x*_1_, (2) it applies a nonlinear function, *f^*^*(*x*), to *x*_1_, (3) it scales *f^*^*(*x*_1_) by a gain factor, *k*_1_, whose magnitude depends on the relative reliability of *t_m_*_1_ and *t_e_*_0_, and (4) it adds *k*_1_*f^*^*(*x*_1_) to *t_e_*_0_ to compute *t_e_*_1_. In the 1-2-Go condition (top), *t_e_*_1_ is the final estimate used for the production of *t_p_*. In the 1-2-3-Go condition (bottom), the updating procedure is repeated to compute a new estimate *t_e_*_2_ by adding *t_e_*_1_ to *k*_2_*f^*^*(*x*_2_) where *x*_2_ is the difference between the second measurement, *t_m_*_2_, and *t_e_*_1_, and *k*_2_ is the scale factor determined by the relative reliability of *t_e_*_1_ and *t_m_*_2_. *t_e_*_2_ is then used as the final estimate for the production of *t_p_*. We assumed that the produced interval, *t_p_*, is perturbed by zero-mean Gaussian noise with standard deviation proportional to the final estimate (*t_e_*_1_ for 1-2-Go and *t_e_*_2_ for 1-2-3-Go) with the constant of proportionality *w_p_*, as in the BLS model. (B) The mapping from measurements to estimates (grayscale) for the EKF estimator in the 1-2-3-Go condition. Red lines indicate combinations of measurements that lead to identical estimates (shown for *t_e_*_2_ = 700, 750, 800, 850, and 900 ms). (C) Mean and standard deviation of *t_p_* as a function of *t_s_* for two example subjects (circles and error bars) along with the corresponding fits of the EKF model (lines). (D) BIAS (circles) and 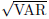 (squares) of each subject (abscissa) and the corresponding values computed from simulations of the fitted EKF model (ordinate). Conventions match Fig 4B. (E) The RMSE in the 1-2-Go and 1-2-3-Go conditions relative to the corresponding predictions from the EKF model (conventions as in Fig 4C). See also S5 Fig.

To further validate the superiority of the EKF model, we directly compared various models to BLS using log likelihood ratio. Specifically, we computed the ratio of the log likelihood of each model given *t_s_* and *t_p_* (*logL*(*M_i_|t_s_, t_p_*) see Materials and methods) to the log likelihood of the BLS model, *logL*(*M_BLS_|t_s_, t_p_*), for each subject. We found that EKF provided the best fit for 8 out of 9 subjects (Table 1). For one of the subjects, the fits were poor for all models but LNE provided the best fit.

**Table 1.** Predictive log likelihood ratio for each model and subject.

| Model | Subject |  |  |  |  |  |  |  |  |
| --- | --- | --- | --- | --- | --- | --- | --- | --- | --- |
|  | CV | GB | LB | PG | SM | SE | TA | VR | VD |
| LNE | 1.1678<br>(0.5012) | -1.8267<br>(0.2976) | -0.3094<br>(0.2020) | 0.1516<br>(0.3407) | **0.4896**<br>**(0.3427)** | -1.7151<br>(0.3493) | -0.0365<br>(0.2806) | 0.3456<br>(0.3593) | 0.3793<br>(0.2175) |
| EKF | **1.1900**<br>**(0.1998)** | **0.0380**<br>**(0.1326)** | **0.7168**<br>**(0.2234)** | **0.3828**<br>**(0.1428)** | -1.4262<br>(0.2755) | **1.0909**<br>**(0.1543)** | **0.2358**<br>**(0.1250)** | **0.3623**<br>**(0.1581)** | **0.6537**<br>**(0.1002)** |
| BLS<sub>mem</sub> | 0.4621<br>(0.1832) | -0.0821<br>(0.0267) | -0.2445<br>(0.0743) | 0.0086<br>(0.0518) | -0.0637<br>(0.2714) | 0.0440<br>(0.0876) | -0.0191<br>(0.0195) | -0.0359<br>(0.0249) | -0.0421<br>(0.0268) |

The predictive log likelihood of each model was calculated as the likelihood of the model given 100 trials of validation data. The model was fit to the remaining data not used for validation. This was iterated until all the data was used for validation (e.g. leave 100 out cross validation). Each entry is the mean of the predictive log likelihood ratio (standard error) between a given model and the BLS model (i.e. $\log[\mathcal{L}(\mathcal{M}_i|t_s, t_p)/\mathcal{L}(\mathcal{M}_{BLS}|t_s, t_p)]$ ). For each subject, the model with the highest log likelihood is shown in bold. See also S6 Fig.

## Discussion

The neural systems implementing sensorimotor transformations must rapidly compute state estimates to effectively implement online control of behavior. Behavioral studies indicate that, at a computational level, state estimation may be described in terms of Bayesian integration [6–15]. However, describing behavior with a Bayesian model does not necessarily indicate that the brain implements these computations by representing probability distributions [21, 22, 57]. Here, we focused on integration of multiple time intervals and found evidence that the brain relies on simpler algorithms that approximate optimal Bayesian inference.

We demonstrated that humans integrate prior knowledge with one or two measurements to improve their performance. A key observation was that the integration was nearly optimal for one measurement but not for two measurements. In particular, when two measurements were provided, subjects systematically exhibited more BIAS toward the mean of the prior than expected from an optimal Bayesian model. This observation motivated us to investigate various algorithms that could lead to similar patterns of behavior.

Analytical and numerical analyses suggested that simple inference algorithms that update certain parameters of the posterior instead of the full distribution can not integrate multiple measurements optimally when the noise is signal-dependent. We then systematically explored simple inference algorithms that could perform sequential updating and account for the behavioral observations. One of the simplest updating algorithms is the Kalman filter [58]. However, this algorithm updates estimates linearly and thus cannot account for the nonlinearities in subjects’ behavior, even for a single measurement (S2 Fig). The LNE model augmented the Kalman filter such that the final estimate was subjected to a point-nonlinearity. This allowed LNE to generate nonlinear biases but since the nonlinearity was applied to the final estimate, LNE failed to capture the decrease in bias observed in the 1-2-3-Go compared to 1-2-Go condition.

Finally, we developed EKF, which is a more sophisticated variant of the Kalman filter that applies a static nonlinearity to the errors in estimation at every stage of updating. This algorithm accounted for optimal behavior in the 1-2-Go condition and exhibited the same patterns of suboptimality observed in humans in the 1-2-3-Go condition. Therefore, EKF provides a good characterization of the algorithm brain uses when there is need to integrate multiple pieces of information presented sequentially. This finding implies that subjects may only rely on the first few moments of a distribution and use nonlinear updating strategy to track those instead of updating the entire posterior. This strategy is simple and in many scenarios could lead to optimal behavior with little computational cost. Moreover, the recursive nature of EKF’s updating strategy allows it to readily generalize to scenarios when it is necessary to update estimates in real time, even when the number of available samples is not known *a priori*, which extensions of our experiment could test.

Maintaining and updating probability distributions is computationally expensive. Moreover, it is not currently known how neural networks might implement such operations [22]. In contrast, EKF is relatively simple to implement. The only requirement is to use the current estimate to predict the next sample, and use a nonlinear function of the error in prediction to update estimates sequentially. Predictive mechanisms that EKF relies on are thought to be an integral part of how brain circuits support perception and sensorimotor function [1, 5, 8, 59–62]. As such, the relative success of EKF may be in part due to its compatibility with predictive coding mechanisms that the brain uses to perform sequential updating. This observation makes the following intriguing prediction: when performing 1-2-3-Go task, subjects do not make two measurements; instead, they use the prior to predict the time of the second flash, use the prediction error to update their estimate, and the new estimate to predict the third flash. In other words, according to the EKF model, the underlying neural signals encode intervals prospectively, which is consistent with electrophysiological experiments in nonhuman primates [63, 64] and rodents [?, ?, ?].

While EKF provides a better account of the observed data in our experiments, it may be that our specific formulation of the Bayesian model did not capture the underlying process. Our BLS model was based on three assumptions: (1) that likelihood function is characterized by signal dependent noise, (2) that the subjective prior matches the experimentally imposed uniform prior distribution, and (3) that the final estimates are derived from the mean of the posterior, which implicitly assumes that subject rely on a quadratic cost function, as was previously demonstrated [24]. Our formulation of the likelihood function is particularly important, as it is the key factor that prohibits simple algorithms such as EKF to optimally integrate multiple measurements. The inherent signal-dependent noise in timing causes the likelihood function to be skewed toward longer intervals (see S1 Appendix). This characteristic feature was particularly important for explaining human behavior in a task requiring interval estimation following several measurements [52]. Moreover, it has been shown that subjects exhibit larger biases for longer intervals within the domain of the prior indicating that the brain has an internal model of this signal-dependent noise [24, 36, 65, 66]. These results support our formulation of the likelihood function. However, one aspect of our formulation that deserves further scrutiny is the assumption that noise perturbing the two measurements was independent. This seems unlikely given the long-range positive autocorrelations in behavioral variability [37, 67, 68], and because S2 is shared between the two measurements, which may lead to correlations between *t_m_*_1_ and *t_m_*_2_.

Our formulation of the prior and cost function should also be further evaluated. For example, humans may not be able to correctly encode a uniform prior probability distribution for interval estimation [34, 36]. Similarly, the cost function may not be quadratic [69]. However, since priors and the cost functions impact both 1-2-Go and 1-2-3-Go conditions, moderate inaccuracies in modeling these components may not be able to explain optimal behavior in 1-2-Go and suboptimal behavior in the 1-2-3-Go condition simultaneously. Finally, recent results suggest the performance may be limited by imperfect integration [70, 71] and imperfect memory [48, 72], which future models of sequential updating should incorporate.

## Materials and methods

### Subjects and apparatus

All experiments were performed with the approval of the Committee on the Use of Humans as Experimental Subjects at MIT after receiving informed consent. Eleven human subjects (6 male and 5 female) between 18 and 33 years of age participated in the interval reproduction experiment. Of the 11 subjects, 10 were naive to the purpose of the study.

Subjects sat in a dark, quiet room at a distance of approximately 50 cm from a display monitor. The display monitor had a refresh rate of 60 Hz, a resolution of 1920 by 1200, and was controlled by a custom software (MWorks; http://mworks-project.org/) on an Apple Macintosh platform.

### Interval reproduction task

Experiment consisted of several 1 hour sessions in which subjects performed an interval reproduction task (Fig 1). The task consisted of two randomly interleaved trial types *−* “1-2-Go” and “1-2-3-Go”. On 1-2-Go trials, two flashes (S1 followed by S2) demarcated a sample interval (*t_s_*) that subjects had to measure [24]. On 1-2-3-Go trials, *t_s_* was presented twice, once demarcated by S1 and S2 flashes, and once by S2 and S3 flashes. For both trial types, subjects had to reproduce *t_s_* immediately after the last flash (S2 for 1-2-Go and S3 for 1-2-3-Go) by pressing a button on a standard Apple keyboard. On all trials, subjects had to initiate their response proactively and without any additional cue (no explicit Go cue was presented). Subjects received graded feedback on their accuracy.

Each trial began with the presentation of a 0.5 deg circular fixation point at the center of a monitor display. The color of the fixation was blue or red for the 1-2-Go and 1-2-3-Go trials, respectively. Subjects were asked to shift their gaze to the fixation point and maintain fixation throughout the trial. Eye movements were not monitored. After a random delay with a uniform hazard (100 ms minimum plus and interval drawn from an exponential distribution with a mean of 300 ms), a warning stimulus and a trial cue were presented. The warning stimulus was a white circle that subtended 1.5 deg and was presented 10 deg to the left of the fixation point. The trial cue consisted of 2 or 3 small rectangles 0.6 deg above the fixation point (subtending 0.2 x 0.4 deg, 0.5 deg apart) for the 1-2-Go and 1-2-3-Go trials, respectively. After a random delay with a uniform hazard (250 ms minimum plus an interval drawn from an exponential distribution with mean of 500 ms), flashes demarcating *t_s_* were presented. Each flash (S1 and S2 for 1-2-Go, and S1, S2 and S3 for 1-2-3-Go) lasted for 6 frames (100 ms) and was presented as an annulus around the fixation point with an inside and outside diameter of 2.5 and 3 deg, respectively (Fig 1A,B). The time between consecutive flashes, which determined *t_s_*, was sampled from a discrete uniform distribution ranging between 600 and 1000 ms with a 5 samples (Fig 1C). To help subjects track the progression of events throughout the trial, after each flash, one rectangle from the trial cue would disappear (starting from the rightmost).

Produced interval (*t_p_*) was measured as the interval between the time of the last flash and the time when the subject pressed a designated key on the keyboard (Fig 1A,B). Subjects received trial-by-trial visual feedback based on the magnitude and sign of the relative error, (*t_p_ − t_s_*)*/t_s_*. A 0.5 deg circle (“analog feedback”) was presented to the right (for error *<* 0) or left (error *>* 0) of the the warning stimulus at a distance that scaled with the magnitude of the error. Additionally, when the error was smaller than a threshold, both the warning stimulus and the analog feedback turned green and a tone denoting “correct” was presented. If the production error was larger than the threshold, the warning stimulus and analog feedback remained white and a tone denoting “incorrect” was presented. The error threshold was scaled with the sample interval to accommodate the scalar variability of timing that leads to more variable production intervals for longer sample intervals (Fig 1D). The scaling factor was initialized at 0.15 at the start of every session and adjusted adaptively using a one-up, one-down scheme that added or subtracted 0.001 to the scaling factor for incorrect or correct responses, respectively. These manipulations ensured that the performance across conditions, subjects, and trials remained approximately at a steady state of 507 correct trials.

To ensure subjects understood the task design, the first session included a number of training blocks. Training blocks were conducted with the supervision of an experimenter. Training trials were arranged in 25 trial blocks. In the first block, the subjects performed the 1-2-Go condition with the sample interval fixed at 600 ms. In the second block, we fixed the interval to be 1000 ms. In the third block, the subject performed the 1-2-3-Go task with the interval fixed at 1000 ms. In the fourth block, the subject continued to perform the 1-2-3-Go task, but with the intervals chosen at random from the experimental distribution. In the final training block, the task condition and sample intervals were fully randomized, as in the main experiment. The subject then performed 400 trials of the main experiment. Subjects completed 10 sessions total, performing 800 trials in each of the remaining 9 experimental sessions. To ensure subjects were adapted to the statistics of the prior [24], we discarded the first 99 trials of each session. We also discarded any trial when the subject responded before S2 (for 1-2-Go) or S3 (for 1-2-3-Go) or 1000 ms after the veridical *t_s_*. S1 Table summarizes the number of completed trials for each subject. Data from two subjects were not included in the analyses because they were not sensitive to the range of sample intervals we tested and their production interval distributions were not significantly different for the longest and shortest sample intervals.

### Models

We considered several models for the interval estimation: (1) an optimal Bayes Least-Squares model (BLS), (2) an optimal Bayes Least-Squares model that allowed different noise levels for the two measurements in 1-2-3-Go trials, (3) an extended Kalman filter model (EKF), and (4) a linear-nonlinear estimation model (LNE). All models were designed to be identical for the 1-2-Go task where only one measurement was available but differed in their prediction for the 1-2-3-Go trials.

### BLS model

We used the Bayesian integration model that was previously shown to capture behavior in the 1-2-Go task [24]. This model assumes that subjects combine the measurements and the prior distribution probabilistically according to the Bayes’ rule:

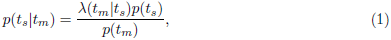

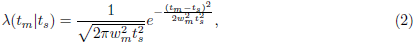

where *p*(*t_s_*) represents the prior distribution of the sample intervals and *p*(*t_m_*) the probability distribution of the measurements. The likelihood function, (*t_m_|t_s_*), was formulated based on the assumption that measurement noise was Gaussian and had zero mean. To incorporate scalar variability into our model, we further assumed that the the standard deviation of noise scales with *t_s_* with constant of proportionality *w_m_* representing the Weber fraction for measurement.

Following previous work [24], we further assumed that subjects’ behavior can be described by a BLS estimator that minimizes the expected squared error, and uses the expected value of the posterior distribution as the optimal estimate:

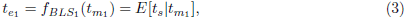

where *f_BLS_*_1_ denotes the BLS function that maps the measurement (*t_m_*_1_) to the Bayesian estimate after one measurement (*t_e_*_1_). The subscript 1 is added to clarify that this equation corresponds to the condition with a single measurement (i.e., 1-2-Go). The notation *E*[*•*] denotes expected value. Given a uniform prior distribution with a range from *t*^min^*_s_* to *t*^max^*_s_*, the BLS estimator can be written as:

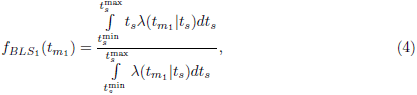

We assume that *t*^min^*_s_* and *t*^max^*_s_* match the minimum and maximum of the experimentally imposed sample interval distribution. We extended this model for the 1-2-3-Go task to two measurements. To do so, we incorporated two likelihood functions in the derivation of the posterior. Assuming that the two measurements are conditionally independent, the posterior can be written as:

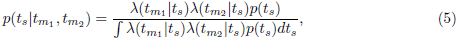

where *t*_m1_ and *t*_*m*2_ denote the first and second measurements, respectively, and the likelihood function, λ, is from Eq (2). Because measurements are taken in a sequence, we can rewrite Eq (5) in a recursive form

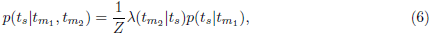

Where *p*(*t_s_|t_m_*_1_) the posterior as specified in Eq (1) and *Z* is a normalization factor equal to the denominator of Eq (5). Although the posterior for Eqs (5) and (6) are identical, specifying the posterior in this way allows for the algorithm to be updated following each measurement.

The corresponding BLS estimator can again be written as the expected value of the posterior:

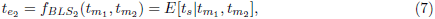

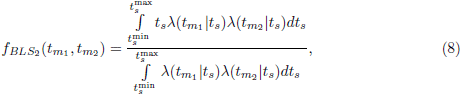

where *f_BLS_*_2_ denotes the BLS function that uses two measurements (*t_m_*_1_ and *t_m_*_2_ to compute *t_e_*_2_. The subscript 2 represents the optimal mapping function for two measurements (i.e., 1-2-3-Go). We performed the integrations for the BLS model numerically using Simpson’s quadrature.

### BLS_mem_ model

We also considered the possibility that the brain may not be able to hold representations of the first measurement or the associated posterior perfectly over time until the time for integration. To model this we assumed two Weber fractions *− w_m_* as formulated in the BLS model and *w_mem_* which adjusts the Weber fraction of the first measurement in 1-2-3-Go trials to account for noisy memory or inference processes. In 1-2-Go trials, the posterior was set according to Eq (1) with *w_m_* controlling the signal dependent noise. In 1-2-3-Go trials, the posterior was set according to

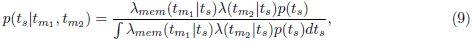

With the likelihood function associated with the first measurement defined as

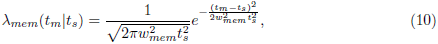

This formulation allows the measurement noise to be different for the two measurements. The optimal estimator was then calculated as

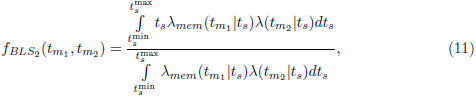

### EKF model

EKF implements an updating algorithm in which after each flash, the observer updates the estimate, *t_e__n_*, based on the previous estimate, *t_e__n−_*_1_, and the current measurement, *t_m__n_*. The updating rule changes *t_e__n−_*_1_ by a nonlinear function of the error between *t_e__n−_*_1_ and *t_m__n_*, which we denote by *x_n_*.

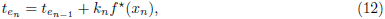

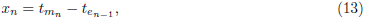

*f^*^* is a nonlinear function based on the BLS estimator, *f*_BLS1_:

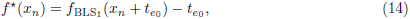

*k_n_* is a gain factor the controls the magnitude of the update and is set by the relative reliability of *t_e__n−_*_1_ and *t_m__n_*, which were formulated in terms of two Weber fractions, *w_n−_*_1_ and *w_m_*, respectively:

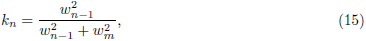

To track the reliability of *t_e__n_* we used a formulation based on optimal cue combination under Gaussian noise. For Gaussian likelihoods, the reliability of the estimate is related to the inverse of the variance of the posterior. Similarly, the reliability of the interval estimate is related to the inverse of the Weber fraction. Therefore, we used the following algorithm to track the Weber fraction of the estimate, *w_n_*:

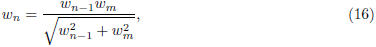

As in the case of Gaussians, this algorithm ensures that *w_n_* decreases with each additional measurement, reflecting the increased reliability of the estimate relative to the measurement. This ensures that the weight of the innovation respects information already integrated into the estimate by previous iterations of the EKF algorithm.

At S1, no measurements are available. Therefore, we set the initial estimate, *t_e_*_0_ to the mean of the prior, and its reliability, *w*_0_, to *∞*. After S2 (one measurement), the EKF estimate is identical to the BLS model. For two measurements, the process is repeated to compute *t_e_*_2_, but the estimate is suboptimal. This formulation can be readily extended to more than two measurements.

### LNE model

LNE uses a linear updating strategy similar to a Kalman filter to update estimates by measurements as follows:

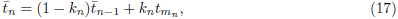

The algorithm is initialized such that ¯*t*_1_ = *t_m_*_1_ and we chose the weighting to be *k_n_* = 1*/n*. This choice minimizes the squared errors in 1-2-3-Go trials. Note that any other choice for *k_n_* would deteriorate LNE’s performance. Following this sequential and linear updating scheme, LNE passes the final estimate through a nonlinear transfer function specified by the BLS model for one measurement (*f_BLS_*_1_):

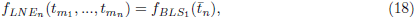

where *f_LNE__n_* denotes the linear-nonlinear estimator after *n* measurements. This formulation ensures that LNE is identical to the BLS in 1-2-Go trials.

### Interval production model

In all models, the final estimate is used for the production phase. Following previous work [24], we assumed that the production of an interval is perturbed by Gaussian noise whose standard deviation scales with the estimated interval. The model was additionally augmented by an offset term to account for stimulus-independent biases observed in responses:

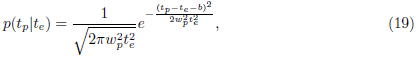

where *w_p_* is the Weber fraction for production, *b* is the offset term, and *t_e_* can refer to the estimate for either 1-2-Go and 1-2-3-Go trials.

All models accommodated “lapse trials” in which the produced interval was outside the mass of the production interval distribution. The lapse trials were modeled as trials in which the production interval was sampled from a fixed uniform distribution, *p*(*t_p_|*lapse), independent of *t_s_*. With this modification, the production interval distribution can be written as:

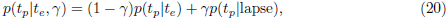

where γ represents the lapse rate. With this formulation, we could identify lapse trials as those for which the likelihood of lapse exceeded the likelihood of a nonlapse. To limit cases of falsely identified lapse trials, we set the width of this uniform distribution conservatively to the range of possible production intervals (between 0 and 2000 ms).

Using simulations, we verified that our model was able to detect lapses for the range of *w_m_*, *w_p_*, and γ values inferred from the behavior of our subject pool. Most subjects had a small probability of a lapse trial that was consistent with previous reports [24]. Two subjects had relatively unreliable performance with a larger number of lapse trials. However, our conclusions do not depend on the inclusion of these two subjects.

### Analysis and model fitting

All analyses were performed using MATLAB R2014b or MATLAB R2017a, The MathWorks, Inc., Natick, Massachusetts, United States. We used a predictive maximum likelihood procedure to fit each model to the data. Assuming that production intervals are conditionally independent across trials, the log likelihood of model parameters can be formulated as:

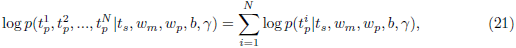

where the superscripts denote trial number. Maximum likelihood fits were derived from N-100 trials and cross validated on the remaining 100 trials. This process was performed iteratively until all the data was fit. The final model parameters were taken as the average of parameter values across all the fits to the data. Fits were robust to changes in the amount of left out data. See S1, S4, and S5 Fig for a summary of maximum likelihood parameters and predictions of each model fit to our subjects.

We evaluated model fits by generating simulated data from that model and comparing various summary statistics (BIAS, 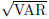, and RMSE) observed for each subject to those generated by model simulations. For the observed data, summary statistics were computed for non-lapse trials and after removing the offset (*b*). Model simulations were performed without the lapse term and after setting the offset to zero. The summary statistics were computed as follows:

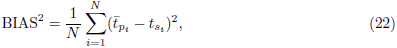

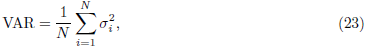

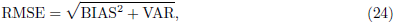

BIAS^2^ and VAR represent the average squared bias and average variance over the *N* distinct *t_s_*’s of the prior distribution. The terms *t_p__i_*, *σ*^2^*_i_* represent the mean and variance of production intervals for the i-th sample interval (*t_s__i_*). The overall RMSE was computed as the square root of the sum of BIAS^2^ and VAR. To find the BIAS^2^ and VAR of each model we took the mean value of each after 1000 simulations of the model with the trial number matched to each subject. This ensured an accurate estimate of these quantities that includes the systematic deviations from the true model behavior due to a finite number of trials.

To perform model comparison, we measured the likelihood of each model, given the maximum likelihood model parameters and the data, *L*(*M_i_|t_s_, t_p_*). We then computed the ratio *L*(*M_i_|t_s_, t_p_*) and *L*(*M_BLS_|t_s_, t_p_*), the likelihood of the BLS model, and computed the logarithm of that value to measure the log likelihood ratio. To generate confidence intervals, we evaluated the likelihoods using 100 trials of test data that were left out of model fitting. We iterated this process until all the data was used as training data, allowing us to measure the variability of the log likelihood ratio for each subject.

## Supporting information

**S1 Fig.**
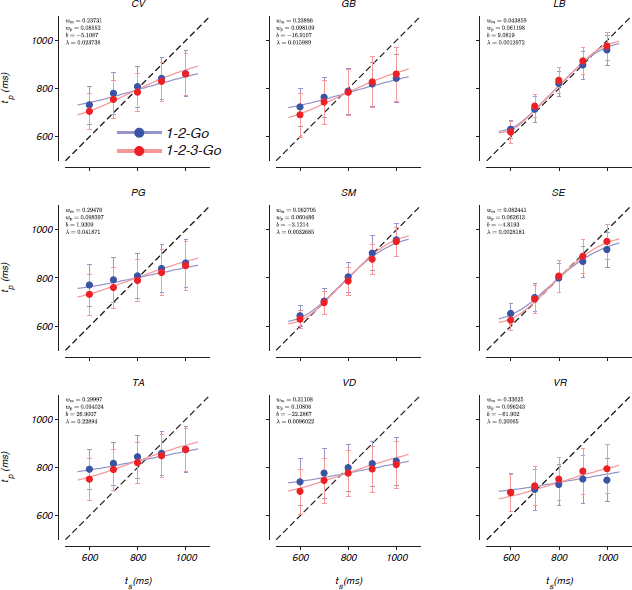
Fit of the BLS model to each subject. Mean and standard deviation (circles and error bars) of the produced interval, *t_p_*, as a function of the sample interval, *t_s_*, for each subject in 1-2-Go (blue) and 1-2-3-Go (red) trials. The dashed line indicates unity and the solid lines depict the BLS model for each subject. Insets: maximum likelihood estimate of model parameters for each subject.

**S2 Fig.**
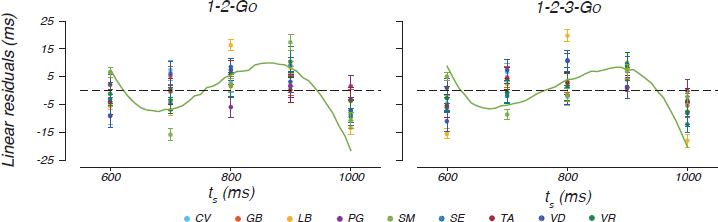
Nonlinear processing of intervals. Mean residuals of the best fitting linear regression model (*t_p_* = *βt_s_* + γ) relating production interval (*t_p_*) to sample interval (*t_s_*) for all subjects (different colors). Each data point represents the mean residual (e.g. 1 *M M Pi*=1 (*t_p__i_ −t_p_*), with *i* indexing trials). The error bars represent the standard error of the residuals. Residuals should be near zero (dashed line) if *t_p_* was linearly related to *t_s_*. The solid green line depicts the expected residuals from the BLS model fits to data from one subject (SM). The predictions for other subjects are qualitatively similar, but the exact shape depends on the measurement noise for each subject. The deviation of the data from the linear model indicates that subjects utilized a nonlinear transfer function to estimate *t_s_*. See also Section 2.4.

**S3 Fig.**
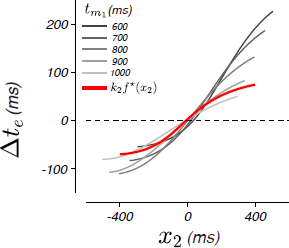
The update function which achieves the BLS after two measurements. The BLS estimator for the 1-2-3-Go task uses the mean of the posterior, *t_e_*_2_ after updating the posterior with the likelihood function associated with the second measurement, *t_m_*_2_. We characterized the function of the BLS estimator in terms of a linear updating term *t_e_*. This term quantified the magnitude by which *t_e_*_1_ has to be updated after the second measurement. A plot of *t_e_* as a function of the error between the first estimate and the second measurement (*x*_2_ = *t_m_*_2_ *− t_e_*_1_) reveals two notables features. First the updating is a nonlinear function of *x*_2_, and second, the form of the nonlinearity depends on *t_m_*_1_ (different grayscales). In other words, BLS can be implemented with a linear updating scheme based on errors only if the update is derived from a nonlinear function whose form of nonlinearity varies with *t_m_*_1_. The red curve corresponds to the nonlinear updating scheme implemented by the EKF model, *k*_2_*f^*^*(*x*_2_). In this case, the updating relies on a nonlinear function of error but the form of the nonlinearity does not vary with *t_m_*_1_. See also Sections 2.3 and 2.5.

**S4 Fig.**
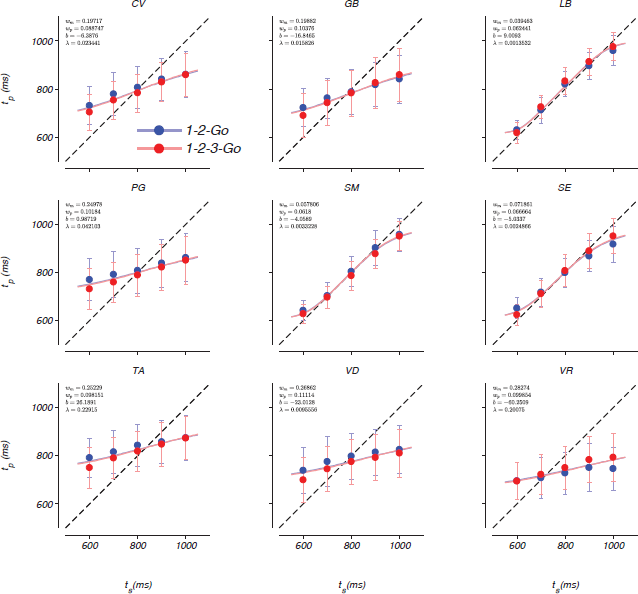
Fit of the LNE model to each subject. Conventions are the same as Supporting information but for the LNE model.

**S5 Fig.**
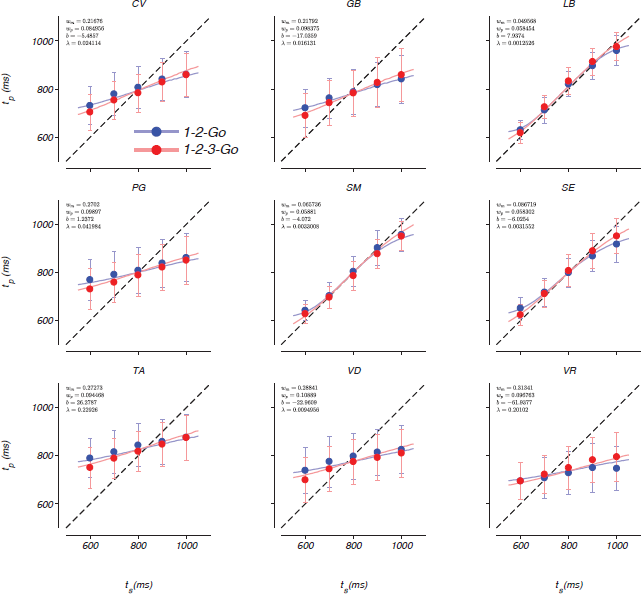
Fit of the EKF model to each subject. Conventions are the same as Supporting information but for the EKF model.

**S6 Fig.**
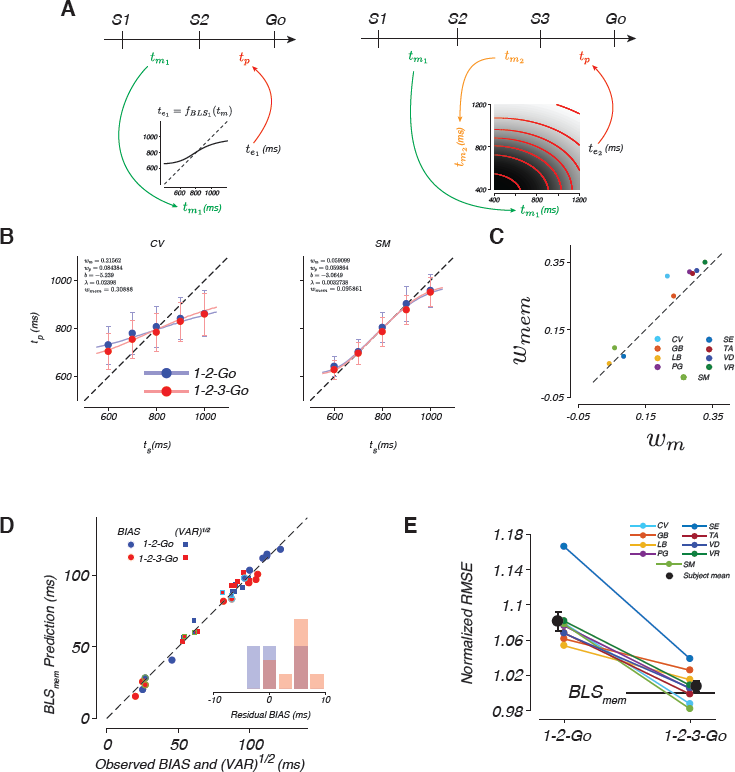
BLS with different levels of noise for the two measurements (BLS_mem_). (A) Left: For the 1-2-Go condition, BLS_mem_ is identical to BLS. Right: For the 1-2-3-Go condition, BLS_mem_ assumes that the Weber fraction for the measurement immediately before production, *w_m_* is the same as the 1-2-Go condition, but the Weber fraction of the first measurement, denoted by *w_mem_*, could be different from *w_m_*. Therefore, the BLS_mem_ model has one more parameter than the BLS model. The bottom grayscale shows the estimate, *t_e_*_2_, as a function of *t_m_*_1_ and *t_m_*_2_ for *w_mem_* = 0.225 and *w_m_* = 0.15. Comparison of the iso-estimate contours (red lines) to the BLS mapping function (Fig 3D), indicates an increased reliance on *t_m_*_2_. (B) Predictions of the BLS_mem_ model (lines) and mean *t_p_* C/- the standard deviation of example subjects (circles and error bars) as a function of *t_s_*. Conventions are the same as panels shown in Supporting information. (C) Maximum likelihood *w_mem_* plot against *w_m_* fit to each subject. Dashed line plots unity. In only one subject *w_m_* exceeded *w_mem_*. (D) Subject BIAS and 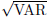 plotted against the predictions of the BLS_mem_ model fit to each subject. Inset: difference between the expected BIAS based on the BLS_mem_ model and that observed. (E) Each subject’s RMSE in the 1-2-Go and 1-2-3-Go conditions to normalized by RMSE expected from the BLS_mem_ model in the 1-2-3-Go condition. See also Table 1.

**S1 Table.** Trials completed by condition and subject in the interval reproduction task. Summary of the number of trials for the interval reproduction task in each condition and the number of trials identified as lapses in parentheses.

|  |  | Subject |  |  |  |  |  |  |  |  |
| --- | --- | --- | --- | --- | --- | --- | --- | --- | --- | --- |
| Trial type |  | CV | GB | LB | PG | SM | SE | TA | VD | VR |
| Total trials | 1-2-Go (lapse) | 3096 (68) | 3156 (36) | 3138 (5) | 3130 (117) | 3120 (10) | 3143 (11) | 2565 (587) | 3099 (18) | 2103 (371) |
|  | 1-2-3-Go (lapse) | 3132 (38) | 3120 (32) | 3169 (1) | 3084 (72) | 3067 (5) | 3159 (3) | 2775 (415) | 3182 (16) | 2108 (316) |

## S1 Appendix

We modeled the likelihood of a sample, *s*, after making a measurement, *m_i_*, perturbed by scalar noise as follows:

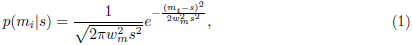

where the parameter *w_m_* scales the standard deviation of the noise with the magnitude of the signal. For *N* conditionally independent measurements with the same *w_m_*, the likelihood can be written as follows:

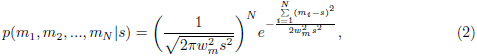

In Eq (2), because the term in the parentheses, which depends on *s*, is exponentiated to the power of *N*, we can conclude that the form of the likelihood function is not invariant with respect to the number of measurements. Therefore, in the presence of scalar noise, to perform exact estimation, it is necessary to update the full distribution after each measurement.

The maximum likelihood estimator (MLE) associated with Eq (2) can be derived analytically by setting the derivative of the logarithm of the likelihood function to zero, which leads to the following solution:

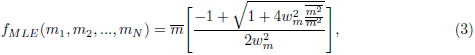

where

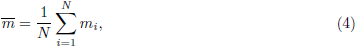

and

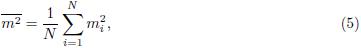

*f_MLE_* is a nonlinear function that depends on the measurements and their squares (e.g. the right side of the equation). Fig 1 demonstrates *f_MLE_* for N = 2. Evidently, MLE for different combinations of measurements is convex. This indicates a larger values of *m_i_* contribute more to *f_MLE_*. This is somewhat counterintuitive because larger *m_i_* is typically associated with larger *s* whose measurement is less reliable. Nonetheless, in the presence of scalar noise, larger measurements contribute more to *f_MLE_*.

**A1 Fig.**
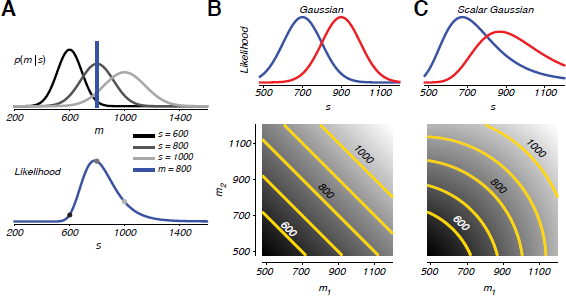
Impact of scalar variability on magnitude estimation. (A) Origin of the skew in the likelihood function for a scalar Gaussian noise process. Top: the distribution of measurements (*m*) for a sample of 600, 800, or 1000 with the Weber fraction for measurement, *w_m_*, set to 0.15. Scalar variability results in a Gaussian of increasing variance for increasing sample intervals. Vertical line plots a hypothetical measurement of 800 which could, with differing likelihoods, come from any of the three distributions. Notice that the large value of *s* is more likely than the small value of *s*. Bottom: The likelihood of each sample interval for a measurement of 800 (blue). Black, dark gray, and light gray circles plot the relative likelihood that the sample interval was 600, 800, or 1000, respectively. Extrapolation from these examples provides intuition for the long tail (skew) in the likelihood of a scalar Gaussian. (B-C) Consequences of skew in the likelihood function for measurement integration. Top row: likelihood functions for two measurements, *m*_1_ = 700 and *m*_2_ = 900 (blue and red, respectively) for (B) Gaussian and (C) scalar Gaussian likelihood functions. Bottom row: mapping functions for maximum likelihood estimators (MLE) corresponding to each likelihood function. Grayscale indicates the magnitude estimate where whiter points indicate greater magnitudes; contours plot loci of equal estimates for different combinations of measurements for estimates from 600 to 1000, spaced by 100. Measurements are given equal weight under the Gaussian noise model which has no skew. This linearity appears in the mapping function as straight contours in the (*m*_1_, *m*_2_) plane (panel B). Skew in the likelihood function leads to a discounting of the smaller measurement in the MLE, as revealed by the curvature of the contours of the associated mapping function (panel C).

## Acknowledgments

We would like to thank Christopher Stawarz for his help in programing the psychophysical paradigm. We also thank Evan Remington, Jing Wang, Hansem Sohn, Devika Narain, and Eghbal Hosseini for their helpful discussions of experimental results. Finally, we would like to thank Joshua Tenenbaum and Evan Remington for helpful comments on an earlier versions of this manuscript, and Rossana Chung for her editorial assistance.

